# RH mapping by sequencing: chromosome-scale assembly of the duck genome

**DOI:** 10.1101/846840

**Authors:** Man Rao, Alain Vignal, Mireille Morisson, Valérie Fillon, Sophie Leroux, Émeline Lhuillier, Diane Esquerré, Olivier Bouchez, Ning Li, Thomas Faraut

## Abstract

Like many other species, the duck genome has been sequenced thanks to the technological breakthrough provided by the emergence of Next Generation Sequencing (NGS). The resulting *de novo* assemblies are however made of thousands of scattered scaffolds. To achieve chromosome-scale contiguity, long-range intermediate genome maps remain indispensable. Radiation Hybrid (RH) maps have been used to assist the generation of chromosome-scale genome assemblies by taking advantage of the high density SNP chips that provide a large number of markers that can be efficiently genotyped on the panel.

In the absence of such a resource in duck, we sequenced 100 hybrid clones of a duck RH panel enabling direct genotyping of the assembly scaffolds on the panel. The rationale is to use scaffolds as markers and to genotype the scaffolds by sequencing the clones: the presence/absence of a scaffold in a particular sequenced hybrid is attested by the presence/absence of reads mapping specifically to this scaffold. The detection of scaffolds exhibiting a chromosomal breakage resulting from the irradiation process revealed itself to be a critical issue of this genotyping by sequencing process. This process resulted in the construction of RH vectors for 2,027 scaffolds, representing a total of about 1 Gb of sequences (95% of the current Duck genome assembly). The subsequent linkage analysis enabled the construction of RH maps and therefore to organize, i.e. order and orient, the scaffolds into pseudomolecules associated to the corresponding duck chromosomes. We describe here the whole mapping process, from sequence-based genotyping to the construction of comparative maps, as well as few examples of intra-chromosomal rearrangements that have been identified by the comparison with the chicken, turkey and zebra finch genomes and subsequently confirmed by FISH.

We describe a method to order and orient sequence scaffolds into super-scaffolds spanning entire chromosomes. The method, which requires a pre-existing RH panel and sequence scaffolds from an NGS assembly, relies on a shallow sequencing of the RH clones. This approach was applied to the duck genome and produced chromosome-scale scaffolds for 29 out of the 41 duck chromosomes.

## Introduction

Two decades after the completion of the first whole genome project of a high eukaryote, namely the human genome closely following the fruit fly genome, thousands of genomes have been sequenced [1] thanks to the emergence of massively parallel sequencing technology. However, the resulting *de novo* genome assemblies remain highly fragmented [2] and only a few dozens of sequenced genomes achieve a reasonable long range contiguity with scaffolds organized in chromosomes [3]. As emphasized by the G10K project and the recent proposed initiative [3] to sequence the genome of all planet’s eukaryotes within the next ten years [1, 3], long range contiguity, ideally chromosome-scale contiguity, remains the final target and is needed to develop in-depth biological analysis that leverage large scale genomic signatures (eg. identification of regulatory elements, chromosome structure and genome evolution). Assembly fragmentation is a direct consequence of the relatively short reads (100-400bp) and small insert libraries (maximum 40kb) used until recently in NGS technology and of the repetitive nature of genomes [4, 5]. The development of methods and technologies to increase the contiguity of genome assemblies is an area of active research [6–8]. On one side of the spectrum, methods that attempt to increase the read length or increase the insert size have been developed. Current single molecule sequencing technologies, such as Oxford Nanopore Technology for example, produce now reads with a median size of 20kb [9] and have been successfully used to increase the contiguity of assemblies [10]. While these approaches significantly improve the length of the resulting contigs and scaffolds the ultimate goal of producing chromosome-scale scaffolds still requires high level ordering information providing long range linkage information between scaffolds. The existing genome assemblies that achieve chromosome scale contiguity, whether hierarchical of whole genome shotgun based, were indeed constructed by combining *de novo* genome assembly and high-density genetic and physical maps, including optical maps, to order and orient sequence scaffolds along chromosomes [5, 7, 11–15]. More recently, new approaches such as the exploitation of high-throughput chromatin conformation structure (HiC) initially developed to study chromosome folding provide information on the one dimensional distance for the that provide have been proposed and applied [6, 8]. This HiC approach is very promising, in contrast to genetic or physical maps, it relies on the wide-spread NGS sequencing technology and can be carried out using the same biological material as the one used for genome sequencing. Recent attempts however underline that local ordering of scaffolds is still problematic and not exempt of errors [6].

RH mapping has been frequently used in assisting the assembly contigs/scaffolds into chromosomes in many species [16–18]. The principle of RH mapping uses the fact that after irradiation of the donor cells, the chromosomes are broken and rescued in recipient cells before being randomly lost during cell culture, and the probability that two markers are co-retained within a same hybrid cell decreases with the physical distance separating them on chromosomes. The RH mapping process requires the genotyping of markers, hundreds or thousands of markers for high-density maps, an intensive and expensive task in the absence of high density SNP arrays [19]. By taking advantage of rapid progress in NGS, we propose here an effective and high throughput method for assigning and ordering contigs/scaffolds onto chromosomes. A duck whole genome RH panel was previously developed [20], in the meantime, duck genome was sequenced using the Illumina Genome Analyzer II technology, providing a de novo genome assembly of 78,487 scaffolds with a N50 scaffold of 1.2Mb [21]. In order to bring this highly fragmented assembly to chromosome-scale contiguity we performed a survey sequencing (0.3X of duck specific sequences) of the 100 clones of the duck RH panel enabling to perform RH mapping using the scaffolds as markers. Using this approach, we were able to organize and order 2,027 scaffolds, covering 95% of assembled duck genome, into 29 duck chromosomes leaving 11 microchromosomes and chromosome W uncovered.

We describe in the following sections the whole mapping process, from sequence-based genotyping to the construction of chromosomal maps. We discuss a few examples of intrachromosomal rearrangements that have been identified by comparison with the chicken, turkey and zebra finch genomes and subsequently confirmed by FISH.

## Results

### Rationale

In a RH panel, the chromosomal fragments resulting from the irradiation of the donor genome are randomly retained in the different clones. The comparison of retention profiles of two markers provides information on their relative one-dimensional distance in the genome. Indeed, two nearby markers behave frequently in a coordinate manner (either co-retained or both absent), because they are likely to be located on the same fragment. The comparative analysis of retention profiles (presence/absence for each clone) is the foundation of RH mapping [19, 22]. In contrast to traditional RH mapping where markers are designed from available genomic sequences, our proposed method treats scaffolds as markers. The presence/absence of a scaffold in a particular sequenced hybrid is attested by the presence/absence of reads mapping specifically to this scaffold. Having in hand an RH panel and composed of 100 clones and the NGS assembly, we sequenced the 100 duck RH hybrid clones enabling to genotype directly the assembly scaffolds. We describe in the following section, the different steps of this mapping by RH sequencing procedure: from the raw sequence data to the retention pattern for the scaffold-markers to the construction of the maps, pseudomolecules and finally a validation of the resulting assembly. Following previous work, we use a comparative mapping approach to order the scaffolds along chromosomes using the chicken genome as reference [23].

### Radiation Hybrid sequencing

The sequencing of the 100 RH hybrid clones produced a total 886 million paired-reads corresponding to 177 Gb of raw sequences. Considering a mean retention of 20% of a haploid duck genome in each hybrid and haploid genome sizes of 1 and 3 Gb for duck and hamster respectively, a hybrid clone contains 6.2 Gb of which 3% is of duck origin (6Gb of hamster and 200 Mb of duck DNA). The expected haploid coverage per hybrid is therefore 0.3 X, for the hamster genome as well as for the duck genome. Out of the 886 million paired reads, 17 million (1.5%) were specifically mapped, in a proper pair, to the duck scaffolds (see Methods) which is close to our expectation of 3% of duck DNA. 12 clones failed to be correctly sequenced leading to a total of 88 clones for subsequent analysis. The amount of sequence produced, and therefore the read coverage, varies considerably between hybrids, (from 2 to 17 million reads, see Supplementary Table 1). Within hybrids, the high variability of read coverage between the different regions of the donor genome is of course the direct consequence of the radiation hybrid construction, missing fragments being exempt of reads (Figure 1). However, even among the fragments that are apparently retained, the coverage variability is higher than the one usually observed when sequencing genomic DNA (Supplementary Figure 2). This coverage variability is consistent with the observed difference in signal intensity of PCR products from RH clones. Both observations argue in favor of a mosaic nature of hybrid cells lines, each clone being in fact a mixture of cells with different duck chromosome complements, their relative proportions explaining the difference in read coverage. An additional reason for this higher variability is the possible retention of only one or both chromosomal segments of the original diploid donor genome (Figure 3). Overall, the variability of the read coverage reflects the fragmented nature of the duck genome in each hybrid cell line, the fragments, resulting from the radiation procedure, being either absent or present but in different proportions. The goal of the calling procedure is to identify those fragments and their status (absence/presence) in each hybrid.

**Figure 1.**
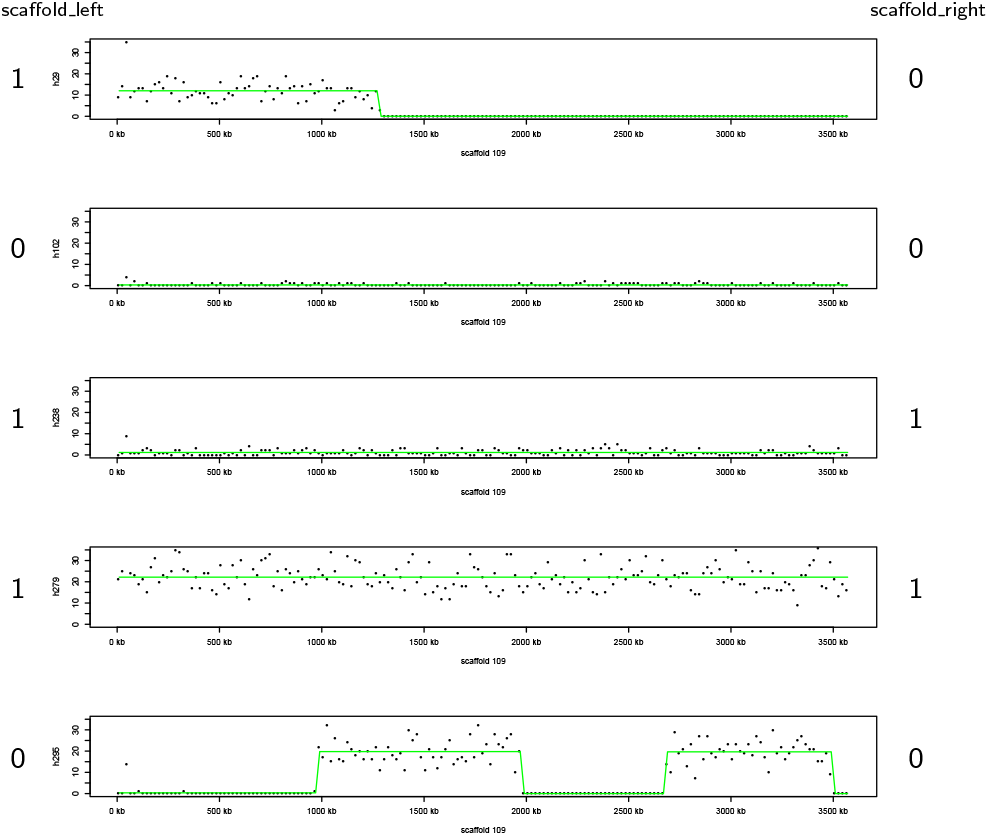
Read depth along a single scaffold. Read depth along a single scaffold in five hybrids. Read counts in 20 kb windows (y-axis) is reported along the scaffold (x-axis) for five RH clones. At least 5 of these fragments can be observed in the hybrid h295 (last line), where read coverage signs the presence absence status. The green lines indicate the mean read count values for the 20 kb windows within segments detected by the CBS. Number (0 or 1) to the left and the right of the figure indicate if the scaffold end is considered absent (0) or present (1) in the RH clone.

### Genotyping scaffolds

The first step in an RH mapping experiment is to genotype the markers, here the scaffolds, i.e to decide if a particular scaffold is present or absent in a particular clone. As expected and exemplified by the segmentation of read coverage on a single large scaffold for different hybrids (Figure 1), many radiation induced breakage occur within scaffolds resulting in portions of the genome fragment covered by a given scaffold as being randomly retained or lost. As a result, and in contrast to RH mapping with locus specific markers, our marker-scaffolds can have different genotypes along their length. However, for our purpose of ordering and orienting scaffolds along chromosomes, where scaffold adjacencies have to be established, the only relevant information is the retention profile at the scaffold extremities. In our genotyping procedure, a scaffold is represented by a couple of markers, one for each extremity, for which a pair of retention profiles have to be determined (Figure 1). Dudchenko *et al.* (2017) used the same idea for their HiC guided assembly by splitting each scaffold in a pair of sister hemi-scaffolds, proximities between scaffolds being derived from proximities estimated between all non-sisters hemi-scaffolds [6]. The problem is now reduced to that of a read coverage segmentation problem in order to determine the retention status, presence or absence, of scaffold extremities (Figure 1). This is essentially a problem of copy number variation detection using sequence data for which many tools have been developed [24]. This apparently simple all or nothing situation (presence/absence) is complicated by different aspects (i) the mosaic nature of hybrid cells lines in which only a variable percentage of cells have retained a donor duck chromosome segment, inducing a sequencing coverage variability that is a nuisance factor (ii) the fragmented nature of the assembly limiting the neighbouring information important to detect shifts in read coverage indicative of a change in copy number variation (iii) the low average read coverage of 0.3X. Following the CNV detection framework using read depth approaches, for each hybrid, the landscape of read counts is segmented into intervals for which the read depth is considered as constant (see Materials and Methods). The RH genotypes (presence or absence) are called on each interval based on the average read counts. When the two extremities of a scaffold exhibited exactly the same pattern of retention, the two corresponding markers were merged into a single marker. For instance, scaffold 109 in hybrid 295 (Figure 1) was segmented into 5 segments by CBS, whereas only the first segment and the last (fifth) segment were used for calling. As a result, 663 scaffolds had at least one RH clone for which the presence/absence genotype was different for each end and had therefore one marker at each extremity, the remaining 1,364 scaffolds were treated as single marker. The total sums up to 2,690 markers covering 1055 Mb of the duck genome assembly.

### Mapping the duck scaffolds on the chicken genome

For the purpose of our comparative mapping approach used to order the scaffolds, they first have to be ordered along the chicken chromosomes according to their sequence similarities with the chicken genome. 1,868 scaffolds (92%) were successfully aligned to the chicken genome (average percent identity 73%, see Supplementary Figure 4). A scaffold was considered successfully aligned if at least 50% of its bases are covered by chains of alignments (see Materials and Methods). Among those scaffolds, 90% had chains covering more than 70% of their length (84% more than 90%). In total, the 2,143 chains covered more than 97% of the assembled chicken genome (the named chromosomes omitting the unplaced contigs) and, except for 41 scaffolds mapping to different chicken chromosomes, it was possible to organize the duck scaffolds into 30 groups, each group corresponding to a single chicken chromosome. Duck scaffolds mapping to different chicken chromosomes are unexpected since they would sign evolutionary interchromosomal rearrangements that are unexpected between chicken and duck [25] and known to be rare between bird genomes [26]. These 41 scaffolds were therefore considered as chimeric.

In order to detect potential intrachromosomal rearrangements between chicken and duck, scaffolds that mapped to different chicken locations (more than 1Mb apart on the same chromosome) were identified as potential signature of chromosomal rearrangements. We identified 126 such breakpoints corresponding to 106 scaffolds (a scaffold can be implied in more than one breakpoint). While this number could be viewed as an underestimation for medium-scale rearrangements because only evolutionary breakpoints that occur within scaffolds are taking into account, since most of the duck genome is covered by the scaffold assembly, we believe that most of rearrangement breakpoints occur within scaffolds. In addition, this number is consistent with what is expected based on the estimated 114 intrachromosomal rearrangements between the chicken and zebra finch genomes that were identified using marker distance of 1 Mb [27]. The same procedure was applied to the muscovy duck genome assembly and resulted in the identification of 153 such breakpoints between chicken and the muscovy genome, corresponding to 103 scaffolds.

### Map construction and Pseudomolecules

Two-point linkage analysis of the 2,690 markers, using a LOD score threshold of 4.5, identified 51 linkage groups. These linkage groups reflect almost exactly the organization of duck scaffolds in chicken chromosomes obtained by sequence similarity and described above (Supplementary Figure 3). One notable exception is the short arm of chicken chromosome 4. This chicken chromosome corresponds to duck chromosomes 6 and 10 and is the only interchromosomal rearrangement between the duck and the chicken genome known to date. The duck chromosome 10, corresponding to the short arm of chromosome 4, is known to harbor the *HPRT* gene used to select for duck chromosomal fragments during the construction of the RH panel. As a consequence the duck scaffolds corresponding to this region have a high retention rate which is too high for RH mapping and were filtered out (see Materials and Methods), explaining the low coverage of scaffold alignments for this chromosomal region.

The agreement between the scaffold composition of the linkage groups and the expected chromosome organization, inferred from alignments of the scaffolds to the chicken genome, provides a first validation of the retention profiles deduced for each scaffold. The chromosomal RH maps of the scaffolds were constructed independently for each chicken chromosome, e.g by merging the linkage groups mapping to the same chicken chromosome, in the exception of chicken chromosome 4 where two separate groups of markers were considered.

The traditional RH mapping approach tries to infer the map order from RH vectors by identifying a map order optimizing a given criteria (minimum obligate breakpoints, multipoint likelihood) [19, 28]. Although this method is suitable for genotyping a small number of markers, it becomes tedious when several hundreds or thousands are involved. In order to build the duck RH maps we used a comparative mapping approach suitable for genome-wide marker ordering, in which a genome phylogenetically close to the genome to be mapped is used as a a prior information for marker ordering [23]. This approach has been used to assist the genome assembly of different animals [18, 29, 30].

In order to estimate the possible influence of the *a priori* information provided by the reference genome on the final duck RH map, we built 3 different duck RH maps using the scaffolds RH vectors for chromosome 2 and the chicken, zebra finch or turkey genome respectively as the reference genome.

The results showed that the maps obtained using the three different species as reference genome were highly consistent (see Figure 2) indicating that the reference genome used had very little impact on the resulting maps. We decided therefore to build the final set of maps using only the chicken genome as the reference order.

**Figure 2.**
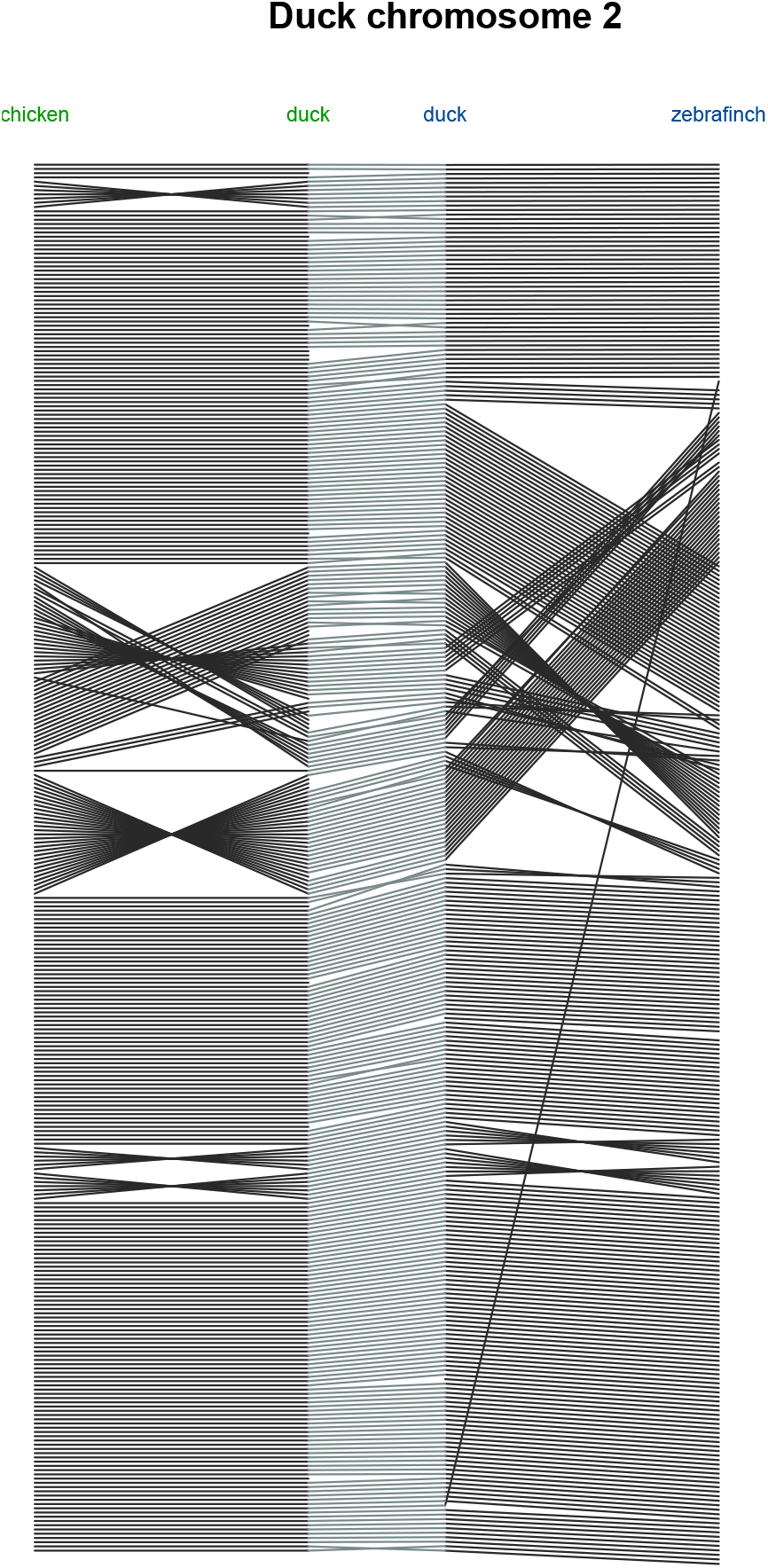
Influence of the reference genome on the RH-based assembly of chromosome 2. Two RH-based assemblies of chromosome 2 are compared, in green an RH assembly constructed with the chicken chromosome 2 as reference and in blue using the zebrafinch as reference underlining that the reference genome as little impact on the reconstructed duck chromosome.

**Figure 3.**
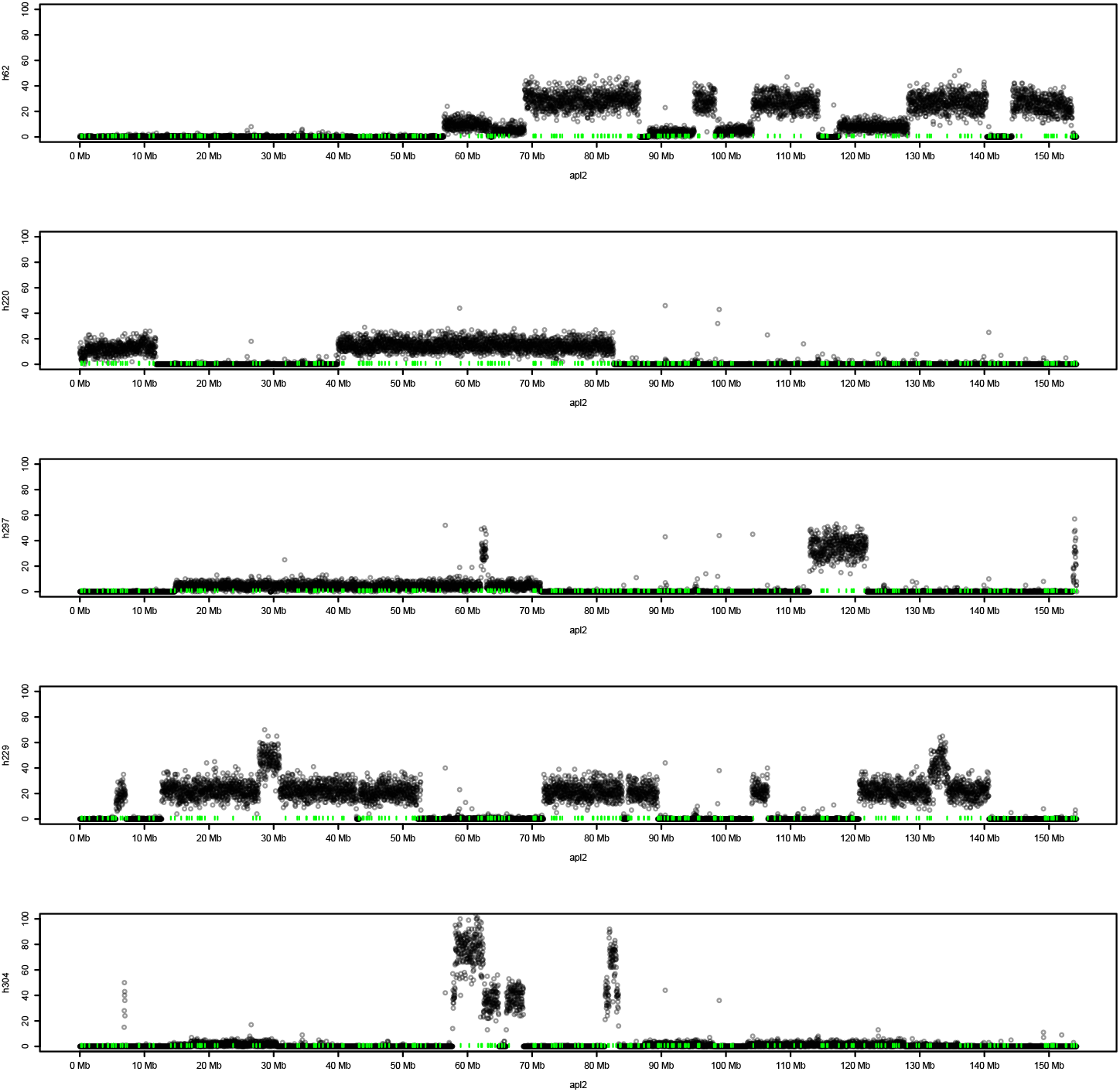
Read depth along the assembled duck chromosome 2. Number of reads per 20kb windows along the assembled chromosome 2 are displayed for five different hybrids. Retained fragments can be clearly observed as as contiguous segments with a constant read depth different from 0 (or background noise). The small vertical green lines represent the scaffolds borders.

In total 29 maps, incorporating 2,343 markers corresponding to 1,657 scaffolds, were built corresponding to duck chromosomes 1-16,18-29 and Z, the map of chromosome 17, corresponding to the chicken 16 chromosome could not be constructed. Among those 1,657 scaffolds, 32 had to be split into parts due to their chimeric nature, that was attested by both the linkage information and by their alignment on the chicken genome. These scaffolds may be considered as putative chimeric scaffolds and the remaining 1,625 scaffolds as non-chimeric scaffolds. Chimeric scaffolds in the duck genome assembly had been already detected in [20]. The resulting maps were used to construct a pseudomolecule for each of the 29 chromosomes (see Materials and Methods). The assembled genome is 970,405,087 bp long representing 89% of the total length of scaffolds.

### Assembly validation using a muscovy duck genome assembly

We used a muscovy duck scaffold assembly (in prep) to check and validate our common duck chromosome assembly. Muscovy and common duck are closely related species having diverged 19 millions years ago [31], in contrast to the evolutionary divergence between duck and chicken which dates back to more than 60 million years [31].

Because it is not the purpose of the current work to assess the correctness of both scaffold assemblies we make the assumptions that the scaffolds are not chimeric and correspond to contiguous regions in both genomes. We therefore focus on the order and orientation, hence adjacencies and relative orientations, of the common duck scaffolds along the chromosomes in our pseudomolecules, as given by our RH maps. The common duck scaffolds we identified as chimeric in our previous analyses were not considered here. In order to detect possible ordering or orientation problems in the proposed common duck chromosomal pseudomolecules, the muscovy scaffolds were aligned on them, which was successful for 1,623 muscovy scaffolds. Because we are interested in medium scale contiguity only muscovy scaffold of size greater than 20kb were considered. Among the 955 muscovy scaffolds satisfying this condition, 791 could be successfully aligned to the common duck assembly (alignment coverage > 0.5 of their length). Two characteristics of our assembly will be tested, the ordering of scaffolds, hence the scaffolds adjacencies defined for each scaffold by its left and right neighbor, and the orientation of scaffolds. The 1,625 scaffolds or our assembly are separated by 1,595 gaps (*n* − 1 − (*k* − 1), where *n* is the number of scaffolds and k the number of chromosomes) that we will try to validate. The second characteristic that will be tested is the orientation of scaffolds. The principle of using two markers per scaffold in order to be able to orient them is limited by the availability of radiation induced breakpoints that enable to separate the two extremities of the scaffold in the radiation mapping context. Single marker scaffolds are therefore difficult to orient and, in addition, even the orientation of two sided markers could be questionable due to the inherent limitation of RH data. We describe next the results of this validation procedure.

### Scaffold ordering

In order to detect possible ordering problems the muscovy scaffolds alignments were chained without strand constraints (see Materials and Methods). Among the 791 muscovy scaffolds under consideration, 757 (95%) could be organized in a single chain suggesting already a good collinearity between the two assemblies. An adjacency between two scaffolds in our assembly is said to be testable if the corresponding gap is locally spanned by a single muscovy scaffold, among the 1,595 adjacencies 1,395 testable adjacencies were identified (87%). A testable adjacency is said to be validated if the order inferred by the muscovy scaffold correspond to the order of the common duck assembly, i.e the alignments surrounding the adjacency gap are adjacent in the muscovy scaffold. Among the 1,395 testable adjacencies 1,346 (96%) correspond to validated adjacencies. Nothing can be said however for the remaining gaps. When the gap is spanned by more than one muscovy scaffold, the nested organization of alignments around the gap is an indication of a potential misassembly. To question the correctness of these problematic adjacencies, a local reordering of the common duck scaffolds was made in order to restore the integrity of the muscovy scaffolds. Because we suppose that the scaffolds are not chimeric, a candidate alternative order of our scaffolds should only reorder common duck scaffolds without splitting them in pieces. Among the 34 muscovy scaffolds organized in more than one chain, the contiguity of only 3 scaffolds could be restored without breaking common duck scaffolds into pieces. We suspect therefore that true evolutionary rearrangements explain most of these discordant orders between the muscovy and the common duck assemblies. Some representative examples of the organization of the pekin assembly along the muscovy scaffolds are given in Supplementary figure 6.

### Scaffold orientation

In order to check the common duck scaffolds orientation within assembly, the muscovy alignments were chained using strand constraints (see Materials and Methods). Possibly wrongly oriented scaffolds were identified on the basis of local discordant orientation along the muscovy scaffold (the orientation of common duck scaffolds inferred by the muscovy chains would imply to reorient a common duck scaffold in our assembly). 28 common duck single marker scaffolds with discordant orientation were identified. Because the orientation of single marker scaffold, is somewhat questionable, we decided to reorient those common duck scaffolds in order to take into account the information provided by the muscovy assembly. It has to be noted that these 28 single marker scaffolds exhibiting a potentially incorrect orientation is far from what would be expected from a process where the orientation of such scaffolds would be decided at random. Indeed, the assembly is composed of 957 single marker scaffolds, if they were oriented at random we would expect half of them being wrongly oriented. The orientation of such single-marker scaffolds were however predicted on the basis of their alignments with the chicken genome: in conserved regions, the common duck genome assembly should exhibit a conserved orientation. 7 additional common duck double marker scaffolds showed incoherent orientation with the muscovy assembly but were left oriented as indicated by the RH maps in order to stay consistent with the whole assembly process.

### The smallest microchromosomes

Among the 2,207 scaffolds for which RH vectors were available, 184 could not be mapped to the chicken genome (mapped means at least 50% could be aligned). As the current chicken genome assembly still lacks the sequence of the 10 smallest microchromosomes, some of these scaffolds could belong to the corresponding duck microchromosomes. We identified 7 linkage groups comprising at least 4 scaffolds using a lod score threshold of at least 4.3. One linkage group covering about 600kb could be assigned to APL17 corresponding to chicken chromosome 16. Another linkage group representing a total of 1.9Mb was identified as homologous to chicken chromosome LGE22C19W28 which is only 950kb long. Another linkage group (how many scaffolds) is homologous to the region 9.9Mb to 19Mb of human chromosome 19 (HSA19) and should therefore correspond to chicken chromosome 30 (Morisson et al, 2007). This result is consistent with our previous result by PCR-based genotyping [20].

### Comparative analysis with chicken

Prior to our study, only little information was available concerning existing chromosomal rearrangements between chicken and duck and most were in the form of limited studies involving FISH mapping of random chicken BAC clones onto duck metaphase spreads [32–34]. Based on our radiation hybrid mapping results, 110 rearrangements between these two species were detected, having precise boundaries. Therefore, in order both to validate the duck RH maps and to confirm the reality of some of the intrachromosomal rearrangements, we tested 4 of them on macrochromosomes by FISH mapping. Chicken BAC clones were selected at the breakpoint boundaries were hybridized onto duck metaphase spreads. All rearrangements between duck and chicken, as detected on the radiation hybrid maps, were confirmed: an inversion on APL1 in the region of 41.5Mb to 51.6Mb; an inversion on APL2 from 66.4Mb to 80Mb; a rearrangement on APL3 and a rearrangement on APL5 (Supplementary Figure 5).

## Discussion

The emergence of high throughput sequencing technologies has enabled the sequencing of a large number of genomes with the objective of producing a reference genome assembly for the corresponding species. While the ultimate goal is always to obtain an assembly with chromosome-scale contiguity, such a long range contiguity remains difficult to achieve. The approaches generally taken nowadays combines long read sequencing technologies together with technologies providing medium to long range distance information, such as optical and chromatin interaction mapping [35, 36]. In most of the cases however, with some notable exceptions [37–39], chromosome scale contiguity is not achieved without the help of other mapping information such as genetic mapping [7, 40, 41]. The challenge lies in the large spectrum of the contiguity information needed, from reads to chromosomes.

While optical maps perform well for medium range distances, it cannot by itself achieve chromosome scale contiguity [37]. On the other hand, although HiC is able to capture chromosome wide distance, it has been reported to be error prone, in particular in the presence of small contigs [6, 36].

Sequencing radiation hybrid clones is presented here as a method for assembling scaffolds to the chromosome level using duck as a test species. Sequencing depth along scaffold segments is used as a test for their presence or absence in a clone. Complete scaffolds or scaffold ends are then used as markers to build radiation hybrid maps and in the latter case, the scaffolds can be oriented. This RH mapping by sequencing approach is combined here with a comparative mapping approach [23] that takes advantage of the existence of a completely sequenced and assembled closely related genome, making the assumption that a limited number of evolutionary chromosomal rearrangements occurred since the divergence of the two species. More precisely, in this context of scaffold ordering problem, the number of chromosomal rearrangements is typically orders of magnitude less than the number of elements to order. As a consequence many of the adjacencies are conserved between the two species. The comparative mapping approach was used here with either the closely related chicken and turkey or the further distant zebra finch genome assemblies as a reference. We observed that the choice of the reference genome has little influence on the resulting maps, providing additional validation of the comparative mapping approach.

In contrast to optical and chromatin interaction mapping, RH mapping requires a RH panel, the construction of which remains labor intensive. Various attempts have been made to simplify or mimic the construction of such a panel, for example the so called Happy mapping approach [42], but have not been conclusive. It is therefore unlikely that an RH mapping like strategy would become the method of choice for ordering scaffolds. Genetic mapping is the natural alternative for long range mapping information but requires the possibility to make crosses. In animals, unless a very large number of individuals are genotyped in a F1 or back-cross design, the resolution will remain low. We would like to stress out, in contrast to strategies involving optical maps or HiC, RH or genetic mapping benefit from the definition of the ordering problem within a solid statistical framework that enables in a natural manner to incorporate additional information such as existing assemblies of closely related species, the basis of our comparative mapping approach.

Different attempts have been made to incorporate comparative genomics information in the assembly process. A closely related assembled genome is used as a reference to assist the assembly if a target species genome making the assumption of a certain degree of synteny conservation. Early attempts have proposed to incorporate comparative information at the very first stages of the assembly process [43–45]. More recent approaches propose to tackle the problem of scaling assembly from scaffolds to chromosomes. The most straightforward approaches propose to organize scaffolds according to their order along the reference genome as predicted by their alignments on this genome [46]. More sophisticated approaches try to combine comparative information with sequence data from the target species [47–49].

With the ever increasing number of genomes that have and are planned to be sequenced in the near future [3] it is likely that comparative approaches will play an important role for the construction of species reference assemblies and the assembly of individual genomes for the characterization of pan-genomes [50, 51].

## Methods

### Common duck and muscovy scaffold assembly statistics

The common duck scaffold assembly (BGI_duck_1., accession GCA_000355885.1) is made of 78,487 scaffolds, 2368 of which are greater than 10kb, with a N50 scaffold size of 1.2Mb and is described in details in [21]. The muscovy duck assembly is composed of 3,702 scaffolds (1,000 scaffolds of length greater than 10kb) with a N50 of 2.4 Mb, having hence a slightly higher contiguity than the common duck scaffold assembly.

### Library preparation and sequencing

The sequencing library was made according to manufacture’s protocol (Illumina). Briefly, 15 μg of genomic DNA was fragmented by sonication and size-selected by separation on agarose gel. Then the fragmented genomic DNA was polished and a single thymine base added to the ends. DNA adaptors with a single thymine base overhang at the 3’end and a 6 nucleotides barcode for multiplexing were ligated to the above products. The mean insert size of the library was 335 bp. One hundred hybrids were sequenced using an Illumina Hiseq2000 sequencing machine. For each hybrid 0.75 pg of DNA was used and twelve hybrids were multiplexed and sequenced in a single lane by pair-end sequencing, with a read length of 101 bases. Individual hybrids are identified by reading the barcode sequence on the adaptors.

### Sequence Alignment and Data Filtering

As the hamster genome sequence was not available at the time of the study, the mouse genome was used as a reference to detect the donor cell sequence reads. Alignment of reads to the mouse genome and to the duck assembled scaffolds were done using BWA [52]. The alignment to the duck genome sequence with BWA was done with the default setting, allowing 6 mismatches per 100 base nucleotides. Only paired reads for which both sequences mapped at unique positions on duck scaffolds were retained for further analysis and reads that could be mapped both on the duck and the mouse genome were discarded. This resulted in an average of 1.6% of sequencing reads per RH clone left for further analysis (Supplementary Table1). After these filtering processes, new bam files were created containing only the paired reads uniquely mapped on duck scaffolds.

### Aligning the duck scaffolds to the chicken genome

All the 19,479 duck scaffolds were aligned to the chicken (Gallus_gallus-4.0, GCA_000002315.2 assembly), zebra finch (taeGut3.2.4 assembly) and turkey (Turkey_2.01, GCA_000146605.1 assembly) genome using the Last software [53]. Large conserved segments are expected between these two closely related species (respectively 80 and 90 million years). However, because of sequence divergence and also of incompleteness of the current assemblies these conserved segments are split into a larger number of local conservation fragments captured by local alignments. In order to recover large conserved segments between the duck scaffolds and the chicken genome we chained the alignments using the chaining algorithm developed in [54]. Briefly, starting with local sequence similarities, conserved segments are chained using an adaptation of the Longest Increasing Subsequence (LIS) algorithm transformed into a general purpose dynamic programming algorithm that identifies high-scoring chain of segments occurring in the same order in both species (see Supplementary information, section 2).

### Scaffold Calling

To detect breakpoints along the scaffolds in the hybrids, the calling was done using the circular binary segmentation (CBS) algorithm introduced by [55], using a window size of 20kb. Fragments or ends of scaffolds not reaching 20kb were not included in the analysis. An output file was generated describing the segmentation of each hybrid, each segment being composed of windows with similar characteristics. This information describes the number of windows in a segment, its first and last window, the total number of reads it contains and the mean value for its 20kb windows. The mean read coverage of a segment was used as a parameter to determine the genotype call: presence or absence of the scaffold segment in the hybrid. Assuming a mean average retention of the duck genome in the RH clones of 20%, an equal distribution of the sequencing depth and a total number of sequencing reads representing 179Gb, for a duck genome fragment present in a clone, there should be on average 3 paired reads aligned per 20kb windows. However, as after data filtering, only 1.6% of the reads were considered as reliably mapped to the duck genome instead of the expected 3% and as the sequencing depth variability was higher than expected (supplementary figure 2), we decided to use the following thresholds: more than 1 read pair per 20kb was considered as presence, lower than 0.5 read per 20kb as absence and in-between as unknown. For each scaffold in each RH clone, we first examined the resulting presence/absence genotype at both ends. If for each of the 86 RH clones the genotype at both ends of a scaffold was the same, the two ends were merged and the scaffold could be treated as a single marker. Contrariwise, if the genotyping result suggested presence of one scaffold end and absence of the other end in at least one of the RH clones, there would be a RH vector for each end and therefore the scaffold could be orientated in the map. To eliminate bad quality markers, we selected only those having a retention higher than 5% in the RH panel and an unknown calling rate less than 15%. To eliminate bad quality markers, we selected only those having a retention higher than 5% in the panel and an unknown calling rate less than 15%.

### Genome maps and chromosome scale assembly

#### Map Construction

Draft maps (comprehensive maps) were made using the comparative mapping approach [23]. Chicken, turkey and zebra finch genomes were used as references to build three sets of maps. First the RH vectors obtained by the scaffold calling and the files containing the ordering of the markers along the reference genomes were used to compute the marker ordering by 2-point likelihoods using the lkh command. Then the properties of the map posterior distributions were obtained with the mcmc command using 32806 as random generator seed and running 5000 mcmc iteration, the first 1000 of which were discarded. The output file from mcmc was used as input for the metamap program described by [56], from which the robust map could be therefore obtained together with the posterior possibility of each maps.

#### Pseudomolecules

The resulting maps were used to construct pseudomolecules for the 29 corresponding duck chromosomes. These maps provide the ordering of scaffolds along the chromosomes. The orientations of scaffold were defined as follows. In the situation where a scaffold is coded as two markers (one at each extremity) and when those two markers are contiguous in the map, the relative order of these two markers in the map provides the orientation of the scaffold. When a scaffold is coded by a single marker, the orientation was defined as the one predicted by their alignment on chicken. The rule being that two consecutive scaffolds with the same orientation are aligned with the same orientation on the chicken genome, that is, on the same strand. For this property to hold, no evolutionary rearrangement must have occurred separating these two regions (scaffold) and modifying their relative orientations but this situation is the exception rather than the rule. We considered therefore that relative strand alignments on chicken of two consecutive duck scaffolds provides the relative orientation of these two scaffolds in the duck assembly and that the orientation of a scaffold in the duck assembly can be predicted by both the orientation of its neighbors and the alignments on chicken.

#### FISH experiments

Nine chicken BAC clones were chosen in the Wageningen BAC library [57] and one BAC in the CHORI-261 chicken BAC library [58] according to their known position, as estimated by BAC end sequence information, in regions paralogous to the breakpoint under study (see table X). BAC clones were grown in LB medium with 5 μg/ml chloramphenicol. The DNA was extracted using the Qiagen plasmid midi kit. FISH was carried out on metaphase spreads obtained from fibroblast cultures of 7-days old chicken and duck embryos, arrested with 0.05 μg/ml colcemid (Sigma) and fixed by standard procedures. The FISH protocol is derived from Yerle et al, 1992, modified for zoo-FISH experiment [33]. Two-colour FISH was performed by labelling 100 ng for each BAC clones with alexa fluorochromes (ChromaTide® Alexa Fluor® 488-5-dUTP, Molecular probes; ChromaTide® Alexa Fluor® 568-5-dUTP, Molecular Probes) by random priming using the Bioprim Kit (Invitrogen). The probes were purified using spin column G50 Illustra (Amersham Biosciences). Probes were ethanol precipitated, resuspend in 50% formamide hybridization buffer (for FISH on chicken metaphases) or in 40% formamide hybridization buffer for heterologous FISH. Probes were hybridised to chicken metaphase slides for 17 hours at 37°C and to duck metaphases for 48H in the Hybridizer (Dako). Chromosomes were counterstained with DAPI in antifade solution (Vectashield with DAPI, Vector). The hybridised metaphases were screened with a Zeiss fluorescence microscope and a minimum of twenty spreads was analysed for each experiment. Spot-bearing metaphases were captured and analysed with a cooled CCD camera using Cytovision software (Leica Biosystem).

## Supplementary information

### Supplementary Tables

**Supplementary Table 1.** Rearrangements confirmed by FISH.

| BAC | Accession | Chicken start (Mb) | Chicken Chr | Duck scaffold | Duck Chr | rearrangement |
| --- | --- | --- | --- | --- | --- | --- |
| BW59P15 | CZ569346 | 42.05 | GGA1p | Sca341 | APL1p | 2*Inversion |
| BW7C7 | CZ560445 | 513 | GGA1p | Sca2083 | APL1p |  |
| BW7I10 | CZ560582 | 67.1 | GGA2q | sca1034_1 | APL2q | 2*Inversion |
| BW23I13 | CZ568657 | 76.5 | GGA2q | sca74_1 | APL2q |  |
| BW17L3 | CZ565516 | 39.97 | GGA3 | Sca1452 | APL3 | 2*Inversion and breakpoint |
| BW18N10 | CZ566381 | 40.53 | GGA3 | Sca1221 | APL3 |  |
| BW66E6 | CZ559900 | 0.395 | GGA5pter | Sca691 | APL5q | small insertion |
| BW8K20 | no | 0.7 | GGA10pter | Sca522 | APL11 | 2*Inversion |
| CH261-179M22 | CC275391.1 | 9 | GGA10 | Sca3697 | APL11 |  |
| BW35L19 | CZ444859.11 | 19 | GGA10qter | Sca428 | APL11 | same location |
| BW19G7 | CZ566048 | 0.4 | GGA11 | Sca2558 | APL12 | 2*Inversion |
| BW20C21 | CZ565661.1 | 7.9 | GGA11 | Sca1176 | APL12 |  |

### Supplementary Figures

**Supplementary Figure 1.**
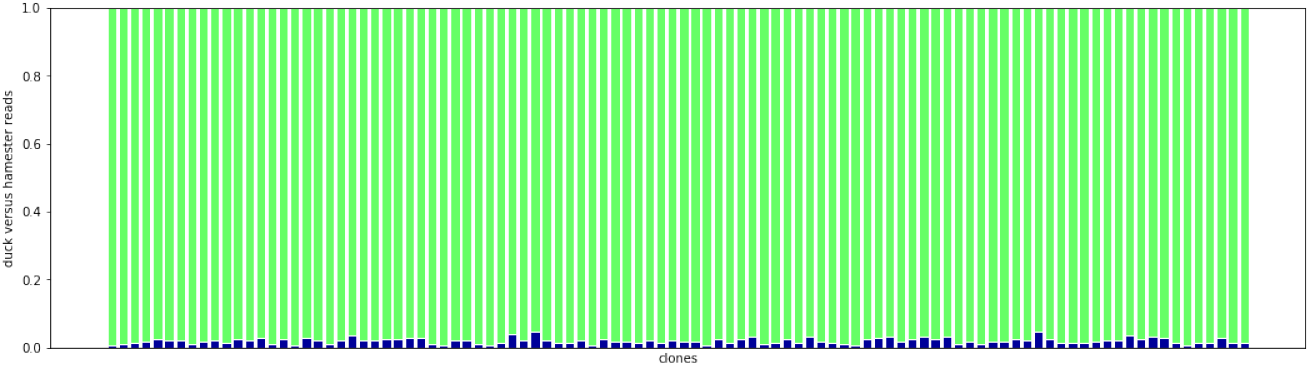
Proportion of useful reads for each clone. For each clone, in blue the proportion of reads that mapped specifically to the duck scaffolds. In total 2.5% of the reads mapped specifically to the duck genome.

**Supplementary Figure 2.**
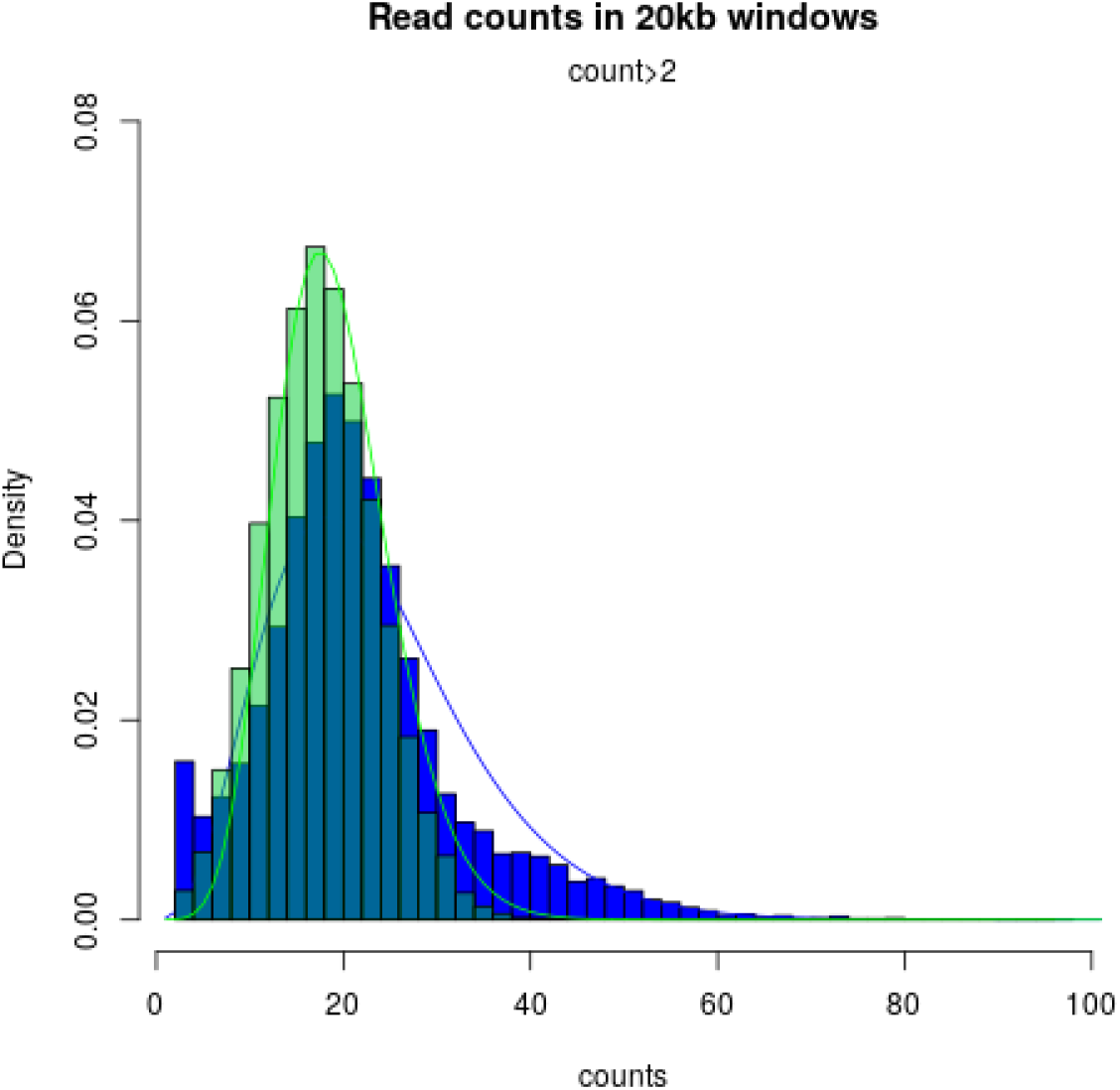
Read depth distribution. Histogram of read counts for 20 kb windows are given for genomic DNA (green) and reads from the donor genome obtained from clones (blue). Lines, green and blue, are the corresponding fitted binomial distributions. The coverage variability is higher than the one usually observed when sequencing genomic DNA.

**Supplementary Figure 3.**
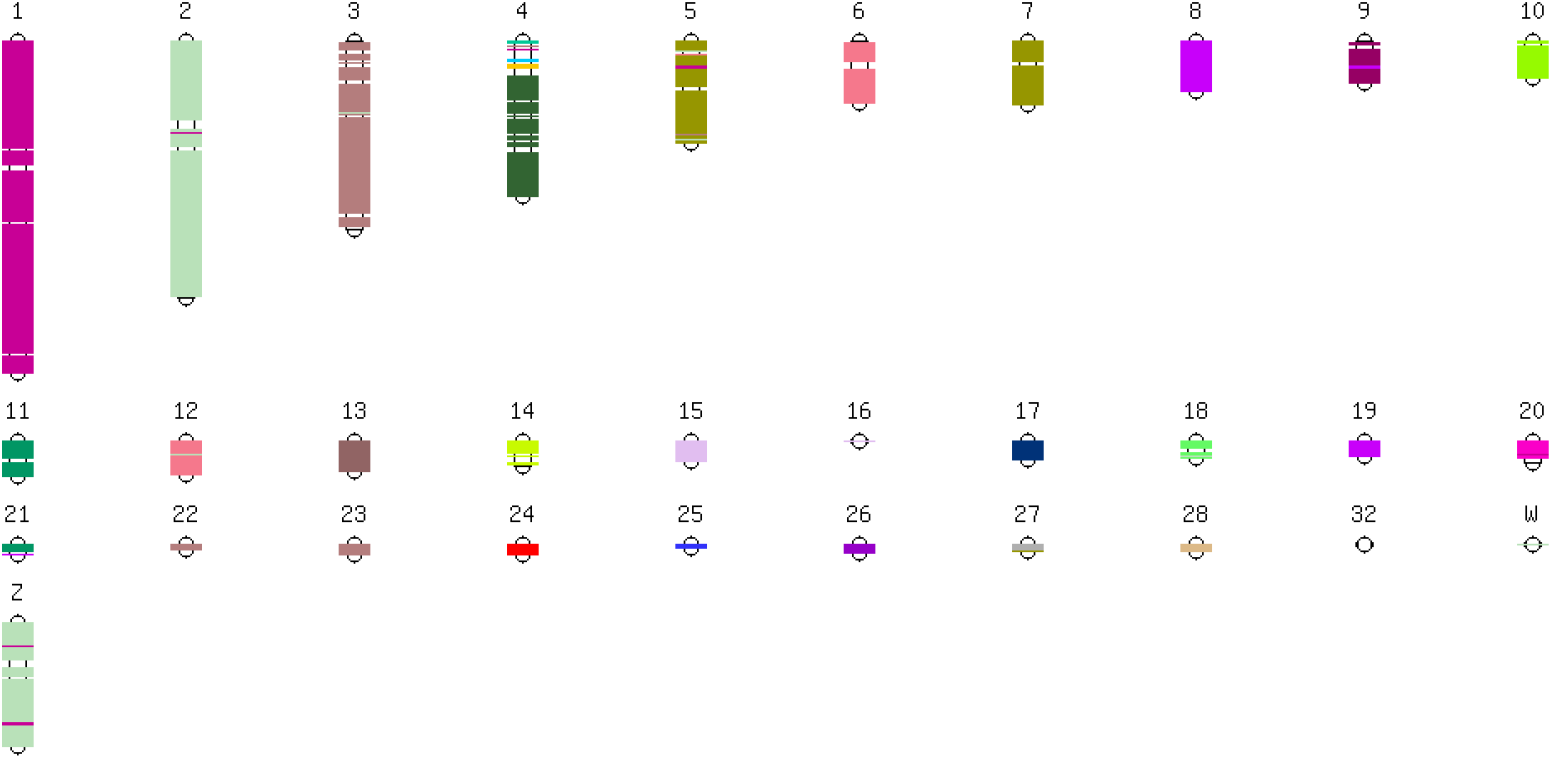
Linkage groups superimposed on the chicken karyotype. Linkage groups superimposed on the chicken karyotype. Duck scafflods are positioned on the chicken genome by sequence alignment whereas colours correspond to duck linkage groups as defined by RH mapping. Apart from chicken chromosome 4 corresponding to two duck linkage groups, as expected based on cytogenetic data, very few inter-chromosomal rearrangements can be detected.

**Supplementary Figure 4.**
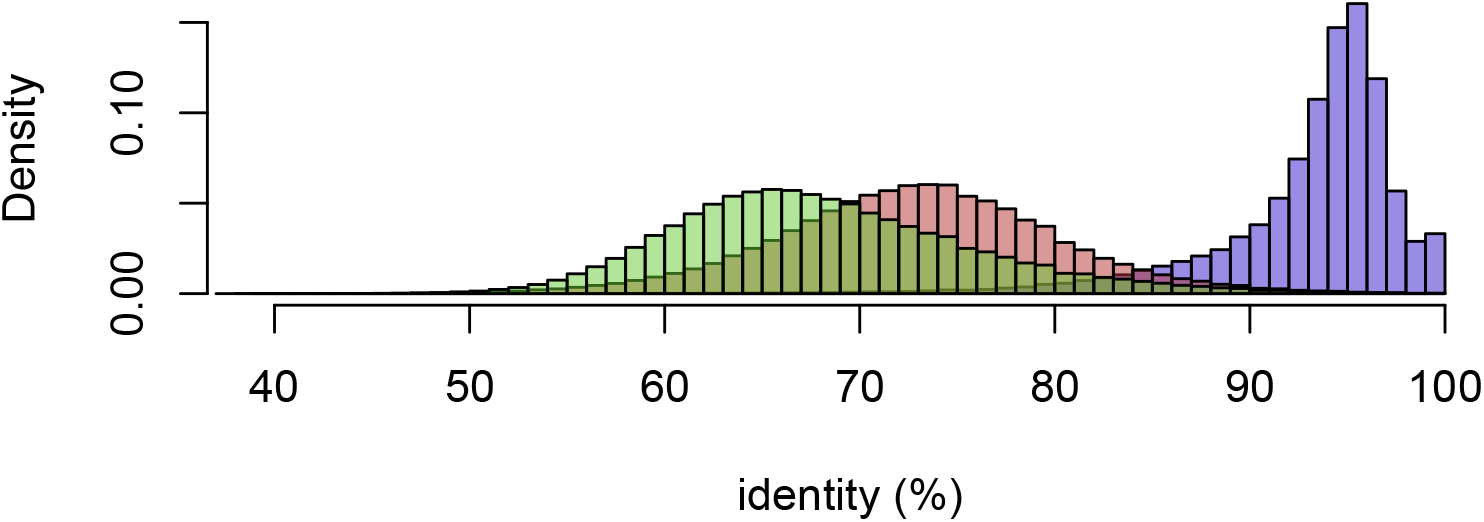
Percent identity. The distribution of percentage identity for the whole genome alignments of pekin genome versus chicken genome, barbarie against pekin and zebrafinch against chicken.

**Supplementary Figure 5.**
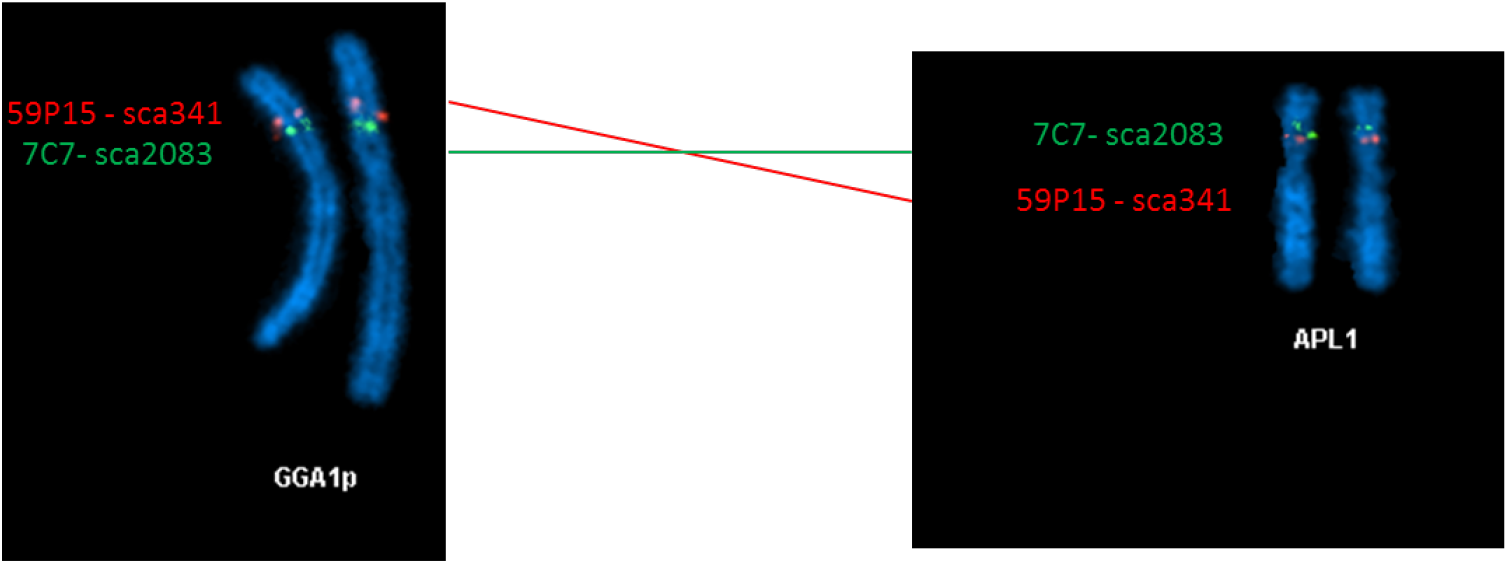
FISH validation. Inversion between the chicken chromosome 1 and Duck chromosome 1 confirmed by FISH.

